# Identification of Engineered IMGT Fc Variants in IMGT/mAb-DB Therapeutic Antibodies and Fusion proteins

**DOI:** 10.1101/2025.09.03.673706

**Authors:** Taciana Manso, Gaoussou Sanou, Christos Nousias, Imene Maalem, François Boutin, Véronique Giudicelli, Patrice Duroux, Marie-Paule Lefranc, Sofia Kossida

## Abstract

Monoclonal antibodies (mAbs) and fusion proteins for immune applications (FPIA) play a crucial role in treating autoimmune diseases and cancers by targeting cell-surface proteins and triggering multiple immune mechanisms. These functions are mediated by the fragment crystallizable (Fc) region of mAbs and fusion proteins, whose interaction with Fc gamma receptors (FcγRs) can be modulated through Fc amino acid (AA) engineering. To address this, we developed the *IMGT/FcVariantsExplorer* tool (https://www.imgt.org/fcvariantsexplorer/) to identify AA changes within the Fc region in mAb and fusion proteins sequences from IMGT/2Dstructure-DB, the AA sequence database of IMGT®, the international ImMunoGeneTics information system®. We used the IMGT® nomenclature of engineered Fc variants involved in antibody effector properties and formats, applying a standardized classification in five categories: ‘Effector’, ‘Half-life’, ‘Physicochemical properties’, ‘Structure’, and ‘Hybrid’. We analyzed sequences of 1,107 mAbs and fusion proteins, identifying 483 entries with Fc AA changes, resulting in 211 unique Fc variants in the dataset. We also used web scraping to retrieve associated biological data from literature. All data have been integrated into IMGT/mAb-DB, with links to sequences in IMGT/2Dstructure-DB, enabling users to query Fc variants by their ‘Category’ or ‘Effect’. This curated dataset reveals key trends in antibody engineering.

## Introduction

Monoclonal antibodies (mAbs) have become a crucial class of therapeutic agents due to their remarkable specificity and efficacy in targeting various diseases (1). Since the landmark approval of the first therapeutic mAb, muromonab-CD3, in 1986 (2), engineered mAbs have undergone significant advancements, including the development of new formats and variants. Their importance in medicine and biotechnology is highlighted by their ability to precisely modulate immune responses, making them essential treatments for a broad range of diseases, from autoimmune disorders to infectious diseases (1).

Immune functions can be modulated by mAbs through their Fragment crystallizable (Fc) region, which interacts with humoral and cellular components of the immune system, such as the C1q complex and Fc gamma receptors (FcγRs) (3,4). These interactions trigger effector functions including complement-dependent cytotoxicity (CDC), antibody-dependent cellular cytotoxicity (ADCC), and antibody-dependent cellular phagocytosis (ADCP). Moreover, the Fc region contributes to prolonging the half-life of mAbs in circulation by binding to the Fc gamma receptor and transporter (FCGRT, neonatal Fc receptor, FcRn) (5).

Currently, all approved mAbs belong to the IgG isotype. This choice is driven by IgG stability, extended half-life, and effector properties (6,7). The selection of IgG1, IgG2 or IgG4 depends on the intended mechanism of action and considers isotypic differences in FcγRs binding and effector properties. IgG2 and IgG4 exhibit fewer interactions with FcγRs than IgG1 (8) and have been used in immune checkpoint inhibitors (9,10). The ability to engineer therapeutic mAbs provides substantial opportunities to modify these interactions with the immune system components and create novel formats (11,12), thereby improving clinical outcomes.

The World Health Organization (WHO) establishes an official nomenclature for mAbs and other therapeutic proteins through the International Nonproprietary Name (INN) Program (13). The biannual INN Proposed and Recommended Lists are highly valuable for therapeutic proteins, providing amino acid (AA) sequences and descriptions that include, for antibodies, the genes and alleles encoding the immunoglobulin chains, and for the constant regions, polymorphic and engineered AA changes (allotypes and variants, respectively). Levering this information, IMGT®, the international ImMunoGeneTics information system® (https://www.imgt.org) (14,15), offers a unique resource for mAbs and fusion proteins with therapeutic applications through two essential databases: IMGT/mAb-DB and IMGT/2Dstructure-DB (16). IMGT/2Dstructure-DB annotates AA sequences from INN/WHO Lists according to IMGT-ONTOLOGY criteria (17,18) and using the IMGT unique numbering for V domains (19) and for C domains (20). IMGT/mAb-DB functions as a repository of therapeutic metadata, providing an intuitive platform for querying mAbs and accessing their annotated sequences in IMGT/2Dstructure-DB. These databases are updated biannually from the INN/WHO lists, while IMGT/mAb-DB is regularly updated with metadata from regulatory agencies, including the U.S. Food and Drug Administration (FDA) and the European Medicines Agency (EMA).

IMGT/mAb-DB originally focused on mAbs (IG), but it now also includes T cell receptors (TR) and other therapeutic proteins, such as fusion proteins for immune application (FPIA), composite proteins for clinical application (CPCA), related proteins of the immune system (RPI). As of July 2025, IMGT/mAb-DB contains 1,769 entries (1,548 IG, 13 TR, 63 FPIA, 75 CPCA and 70 RPI) of which 1,586 have an INN, including the latest updates from INN/WHO List PL132 (2025). Of these, 1,482 entries have their AA sequences fully annotated in IMGT/2Dstructure-DB.

In this study, we focus on identifying engineered AA changes in the constant region of mAbs and FPIAs present in IMGT/2Dstructure-DB. AA changes can be visualised in the IMGT/2Dstructure-DB chain description (per domain, where they are highlighted in orange) or identified using the IMGT/DomainGapAlign tool (16). For standardization, comparisons are made relative to the closest gene or isotype. Using this approach, the IMGT® Nomenclature of engineered Fc variants involved in antibody effector properties and half-life of therapeutic antibodies has been characterized (11,12). Preliminary analyses of engineered variants in INN entries have been performed (21–23). To facilitate such investigations, an automated tool called *IMGT/FcVariantsExplorer* has been developed, enabling rapid and accurate analysis of these AA changes and enhancing data curation. Additionally, we examine the biological significance of these amino acid changes through an extensive literature web-scrapping process.

All the data analyzed in this study are available in IMGT/mAb-DB (https://www.imgt.org/mAb-DB/), where mAbs can be queried based on their variant categories: ‘Effector’, ‘Half-life’, ‘Physiochemical properties’, ‘Structure’, and ‘Hybrid’, offering a valuable resource for evaluating and categorizing engineered mAbs.

## Results

### IMGT engineered variants in IMGT/mAb-DB and IMGT/2Dstructure-DB

In this study, we analyzed the AA sequences of 1,482 INN entries from IMGT/mAb-DB and IMGT/2Dstructure-DB. Among these, 1,310 contained at least one constant domain, and 655 entries carried modifications in the Fc region. To identify and characterize AA changes within Fc regions, we used the IMGT/FcVariantsExplorer tool in combination with an *in-house* text mining tool targeting relevant scientific literature. The identified AA changes were classified according to the IMGT® nomenclature for engineered variants, detailing the AA changes together with their properties and functions. This approach enabled the characterization of 611 engineered molecules. In total, 251 unique variants were identified in the dataset, primarily in human antibodies. Among these, 44 entries contained AA changes for which no functional information could be retrieved from the available literature and therefore were classified as having ‘unknown effects’ (Supplementary Table 1).

Although this article focuses on human therapeutic antibodies, veterinary mAbs are also present in IMGT/mAb-DB, with 20 modified mAb entries from dog, cat, and mouse antibodies. We identified nine Fc variants described in the literature, while 14 remain classified as having “unknown effects”. These veterinary data are not discussed further here; however, all human and veterinary data are publicly available in IMGT/mAb-DB and can be queried using dedicated fields, such as ‘Category’ (e.g., ‘*Effector*’, ‘*Half-life*’) or ‘Effects’ (e.g., ‘*ADCC reduction*’ or ‘*Half-life increase*’). This allows efficient identification and comparison of antibodies carrying specific AA changes. For instance, to retrieve a subset of mAbs designed with Fc silencing, a common strategy in autoimmune disease therapy to minimize Fc-mediated effector functions, users can select “ADCC and CDC reduction” in the ‘Effect’ field and refine the query by choosing a ‘Clinical domain’ or ‘Clinical indication’ in the database, such as ‘Immunology’. This combined search strategy allows targeted access to mAbs specifically designed for autoimmune applications (Figure 1).

**Figure 1.**
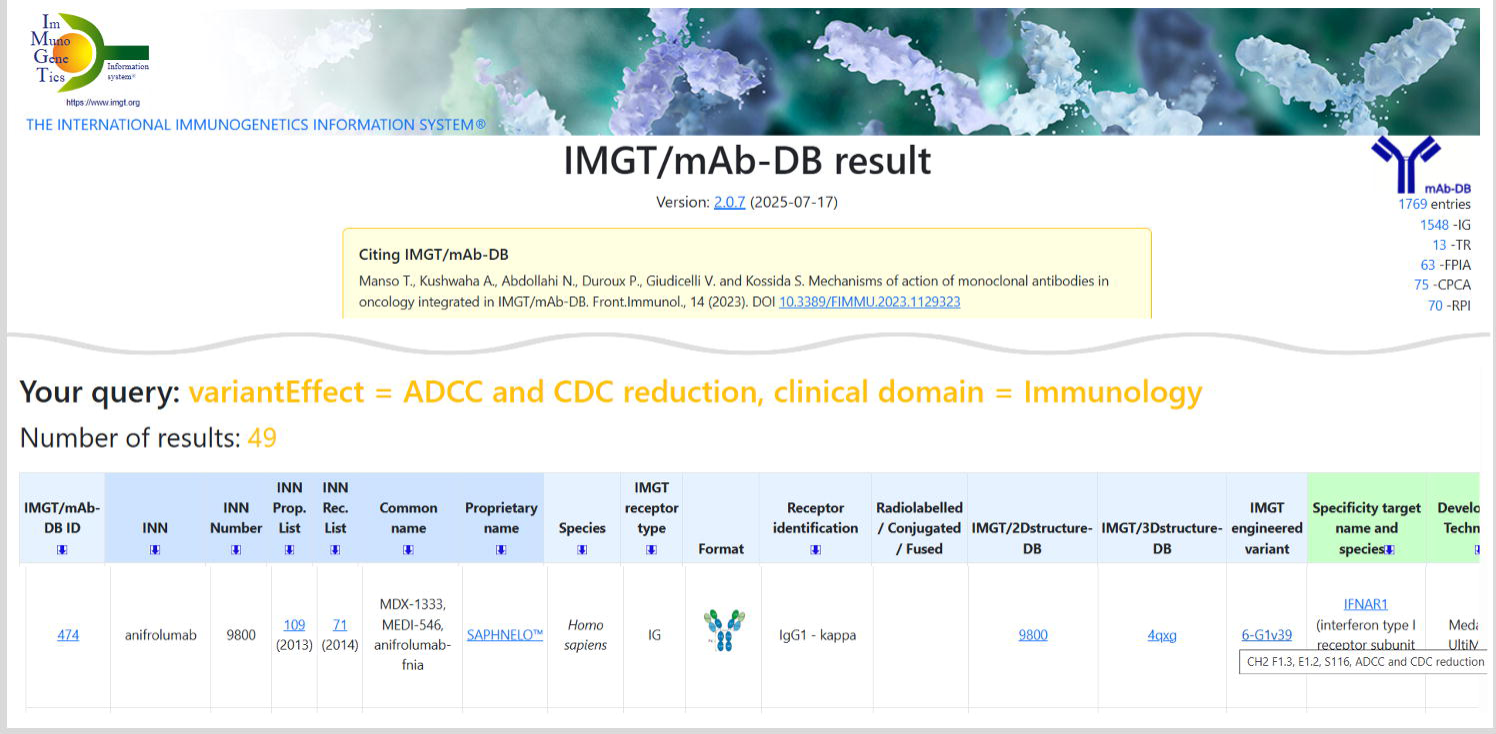
Extract from the IMGT/mAb-DB results page illustrating a query to retrieve mAbs or fusion proteins engineered to silence Fc effector properties. The search combines the filters ‘Effect: ADCC and CDC reduction’ and ‘Clinical domain: Immunology’, resulting in the identification of 49 molecules specifically designed to minimize Fc-mediated effector activity in the context of autoimmune diseases.

### Classification of IMGT engineered variants

Designing therapeutic antibodies requires careful consideration of immunoglobulin isotype variations, including their structural characteristics, half-life, and receptor-binding properties, all of which influence clinical efficacy. Although alternative isotypes in our dataset, such as IgA (one entry), IgE (one entry), and IgM (two entries), display therapeutic potential, their development remains limited due to structural complexity, short half-life, and manufacturing challenges. By contrast, IgG subclasses remain the preferred format, owing to their favorable pharmacokinetics, robust production processes, and well-characterized Fc-mediated effector functions.

The therapeutic mAbs in the dataset are predominantly IgG1 (74%), which is known for its potent effector functions (ADCC and CDC). IgG4, accounting for 18%, is selected for therapeutic applications requiring reduced Fc effector activity. IgG2, comprising 6.5%, is valuable for its stability and antigen cross-linking ability, whereas IgG3 (one entry) is rarely used due to its short half-life and increased susceptibility to proteolysis, despite potent effector functions (6).

The IMGT® engineered variants nomenclature (11) incorporates isotype information to categorize AA changes and their properties and functions. Each isotype is associated with a distinct set of variants, even when AA changes overlap across genes. For instance, variants 6-G1v14 and 6-G4v4 share identical CH2 domain AA changes (A1.3, A1.2) but they are considered distinct due to their occurrence in different genes. Literature evidence indicates that both variants enhance ADCC and CDC within their respective isotype, emphasizing similar functional outcomes despite isotype differences. Functional effects are not generalized across isotypes without experimental support. For a comprehensive list of IMGT nomenclature for engineered variants, see the IMGT Biotechnology page (https://www.imgt.org/IMGTbiotechnology/IGHG_variant/Tableau1.html).

A single mAb chain may include multiple Fc variants, either within the same category or spanning different categories, to optimize its therapeutic efficacy. For instance, *riliprubart* (IMGT/mAb-DB ID: mAbID <u>1426</u>) carries three variants: 6-G4v3 (CH2 E1.2) to reduce ADCC and CDC, 9-G4v24 (CH3 L107, S114) to extend half-life, and 12-G4v5 (h P10) to stabilize hinge and prevent half-IG exchange. Consequently, when variants are analyzed, the same mAb may be represented in multiple groups.

Overall, most entries with engineered Fc regions were classified in the ‘Structure’ category (379 entries), mainly driven by IgG4 antibodies carrying the hinge stabilization variant 12-G4v5 (h P10), comprising 211 entries. The ‘Effector’ category is also highly represented (336 entries), reflecting the widespread use of Fc engineering to modulate antibody effector activities. In comparison, fewer entries are assigned to ‘Half-life’ (83 entries), ‘Physicochemical properties’ (36 entries), and ‘Hybrid’ (12 entries), the latter typically combining constant domains from different isotypes to further modulate Fc functions (Figure 2).

**Figure 2.**
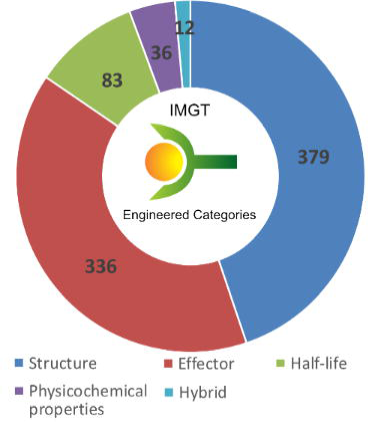
Distribution of Fc-engineered monoclonal antibodies across the five IMGT variant categories: ‘Effector’, ‘Half-life’, ‘Physicochemical properties’, ‘Structure’, and ‘Hybrid’. The counts represent the number of molecules carrying at least one Fc variant within each category. As individual antibodies may include multiple Fc variants, a single molecule can be represented in more than one category.

### ‘Effector’ category

In our dataset, 117 engineered variants are associated with modifications to effector function. Of these, 86 are designed to silence Fc-mediated activity, 22 to enhance it, three variants exhibited dual effects (enhancing ADCC while minimizing CDC or *vice-versa*), and six targeted B cell inhibition. Integrating multiple variants within a single mAb allows precise modulation of Fc function. For example, *evorpacept* (mAbID <u>1293</u>) combines 6-G1v14-1 (CH2 A1.3, A1.2, A1) with the aglycosylation variant 8-G1v29 (CH2 A84.4) to effectively eliminate both ADCC and CDC. Similarly, *nivatrotamab* (mAbID <u>1138</u>) employs 5-G1v20 (CH2 A105) to reduce CDC and 8-G1v29 (CH2 A84.4) to abolish ADCC.

Overall, 336 antibodies and fusion proteins carry engineered Fc regions to alter effector functions. Of these, 310 (approximately 92%) were modified to silence Fc-mediated activity, whereas only 24 (about 7%) were engineered to enhance effector functions. Notably, *botensilimab* (mAbID <u>1123</u>) incorporates the 2,5-G1v8 variant (CH2 D3, L115, E117) to enhance ADCC while minimizing CDC, reducing C1q binding and preventing complement activation. Consequently, its immune modulation relies primarily on effector cells through ADCC, rather than the complement pathway, which can be associated with side effects (24).

‘Effector’ variants are predominantly from the IgG1 subclass, representing 266 entries (80%), followed by IgG4 with 57 entries (17%) and IgG2 with 17 entries (3%). Figure 3 highlights the three most frequent variants for each isotype. In the IgG1, 6-G1v14 (CH2 A1.3, A1.2), commonly known as L234A/L235A or LALA, was the most prevalent, present in 55 entries (21%). This variant reduces recruitment of immune effector cells and suppresses complement activation by decreasing binding affinity to FcγRIIIa and C1q, minimizing unwanted immune responses (25). The second most utilized variant, aglycosylation 8-G1v29 (CH2 A84.4), occurred in 31 entries (12%), removing the glycan at CH2 N84.4 and disrupting FcγRs binding to eliminate effector functions (26). The third most common variant, 6-G1v14-1 (CH2 A1.3, A1.2, A1), observed in 22 entries (8%), extends the LALA mutation with G1>A (G237A) mutation in the CH2 domain, further reducing affinity for FcγRI and FcγRIII and enhancing thermal stability (27).

**Figure 3.**
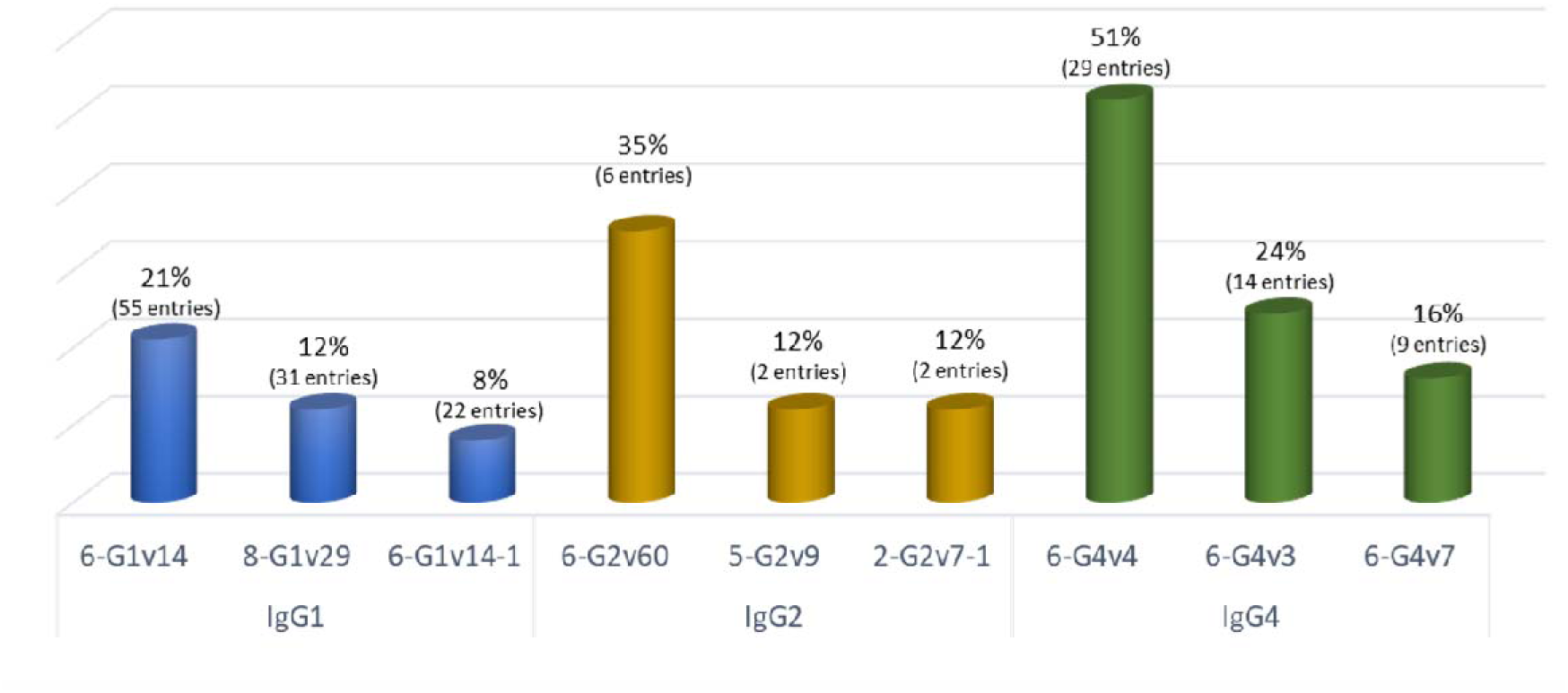
Frequency of the three most common ‘Effector’ Fc variants across IgG subclasses. IgG1 variants are represented in blue, IgG2 in yellow, and IgG4 in green.

Although IgG4 and IgG2 naturally exhibit low FcγRs affinity and limited effector functions (8), specific AA changes have been introduced to further reduce FcγR binding. In IgG4, the most common variants include 6-G4v4 (CH2 A1.3, A1.2), present in 30 entries (52%); 6-G4v3 (CH2 E1.2), found in 14 entries (24%); and 6-G4v7 (CH2 P1.4, V1.3, A1.2, delG1.1), observed in 9 entries (16%). In IgG2, 6-G2v60 (CH2 S115, S116), appearing in six entries (35%) and 6-G2v3 (CH2 A1.2, A1, S2, A30, L92, S115, S116), found in two entries (12%), were most frequent. These engineered modifications strategically abolish unwanted effector functions, enhancing therapeutic efficacy and safety (28).

In contrast, the IgG2 variant 2-G2v7-1 (CH2 D3, A110, E117), observed in two entries, enhances ADCC against the target cell, overcoming IgG2’s naturally weaker FcγRIIIa binding boosting its ability to drive strong immune responses, particularly ADCC (8). This modification makes IgG2-based therapeutics more effective in clinical applications that require robust ADCC. *Tafasitamab* (mAbID <u>522</u>) and *talacotuzumab* (mAbID <u>754</u>), which are IgG1-IgG2 hybrid mAbs (discussed below), incorporate 2-G2v7-1 to enhance ADCC, thereby improving therapeutic efficacy. Apart from these hybrids, all mAbs engineered for enhanced effector function are IgG1 (24 entries), including *enoblituzumab* (mAbID <u>590</u>) and *margetuximab* (mAbID <u>473</u>), which carry modifications to increase effector properties.

Additionally, production strategies such as afucosylation, enzymatic removal of fucose from Fc glycans, enhance ADCC by increasing FcγRIII binding (29). Although our pipeline identifies only sequence-based modifications, afucosylated mAbs can be queried in IMGT/mAb-DB via the ‘Development technology’ field, providing a complementary resource for investigating this enhancement strategy.

### ‘Half-life’ category

Another category of Fc variant targets the antibody interaction with the neonatal Fc receptor (FCGRT, FcRn), which plays a critical role in antibody recycling and serum half-life. Optimizing antibody half-life is crucial for improving therapeutic efficacy and reducing dosing frequency (30). Extending the half-life can be achieved by incorporating substances into the antibody construct, such as PEGylation or albumin fusion. However, advances in Fc engineering allow precise modulation through targeted sequence modifications (31). In total, 25 engineered variants in our dataset are related to half-life modulation.

Most antibodies and fusion proteins carry modifications that enhance FcRn binding and prolong half-life while maintaining pH-dependency release into the serum. Nineteen Fc variants designed to enhance FcRn interaction were identified, encompassing 76 entries. These mutations improve binding at acidic pH in endosomes while allowing efficient release at neutral pH. In IgG1, the most frequent variant was 9-G1v21 (CH2 Y15.1, T16, E18), present in 28 entries (34%). Similarly, in IgG4, the 9-G4v21 variant (CH2 Y15.1, T16, E18) was the most common, observed in seven entries (8%).

In some therapeutic applications, particularly in autoimmune diseases, reducing endogenous antibody half-life by blocking FcRn interactions is advantageous. Seven mAbs in our dataset, including *faricimab* (mAbID <u>793</u>) for ocular treatments (32,33) and *tuvonralimab* (mAbID <u>1204</u>), an anti-CTLA4 antibody, are engineered to reduce FcRn binding, corresponding to variants 9-G1v77-1 (CH2 A15.2, A93, CH3 A115) and 9-G1v77-2 (CH2 K17), respectively, for shortening their half-life (34). *Efgartigimod alfa* (mAbID <u>731</u>), a Fc-gamma fragment, employs ABDEG^TM^ technology (35) with the Fc-engineered region 9-G1v22-1, exhibiting high-affinity and pH-independent binding to FcRn. This design acts as an FcRn antagonist, continuously blocking FcRn interaction with endogenous antibodies and thereby reducing their half-life.

### ‘Structure’ category

The ‘Structure’ category encompasses Fc variants that modify the structural features of mAbs and FPIAs, improving stability, integrity, and functional versatility. These modifications address critical challenges, including promoting heterodimerization for bispecific antibodies, enabling site-specific drug conjugation in antibody-drug conjugates (ADCs), preventing half-IgG exchange in IgG4, and facilitating advanced designs such as IgG hexamerisation for enhanced complement activation.

Human IgG4 can naturally form bispecific antibodies through a dynamic process known as “half-IgG exchange” or “Fab-arm exchange”, in which a half-IgG from one molecule swaps with another (36). To prevent this in therapeutic IgG4 antibodies, stabilizing AA changes are introduced. The hinge variant 12-G4v5 (h P10) stabilizes the structure and prevents half-IG formation (36). Additional CH3 domain modification (12-G4v6 CH3 K88) further improve stability (37). In our dataset of 234 therapeutic IgG4 mAbs, 211 incorporate at least one of these stabilizing AA changes. Notably, half-IG exchange mechanism has also been repurposed to create stable bispecific IgG1 molecules, a concept discussed further below.

To enhance antibody stability, covalent linkages, such as engineered disulfide bonds, have been introduced. Aglycosylated antibodies which lacking the glycan moieties, additional disulfide bridges maintain the structural integrity of CH2 domains (e.g., 11-G1v54 CH2 C83, C85), preserving the antibody native conformation. In the dataset, 12 aglycosylated mAbs, including *acapatamab* (mAbID <u>1074</u>) and *emerfetamab* (mAbID <u>1055</u>), incorporate additional CH2 disulfide bridges.

Fc modifications are crucial for designing bispecific antibodies (bsAbs) which require correct heavy chains (H-H) heterodimerization. Strategies to facilitate efficient heterodimerization often involve modifications to both heavy chains to enhance their interaction. The knob-into-holes approach introduces specific mutations in the CH3 domain, a “knob” (larger amino acids) on one chain and a “hole” (smaller amino acids) on the other, to promote correct pairing (38–40). Four Fc variants employ this approach: 14-G1v26, 14-G1v31, 14-G1v32, and 14-G1v33. For example, 14-G1v26 (CH3 Y22) pairs with 14-G1v31 (CH3 T86) on the other chain.

Another technique for achieving heterodimerization involves electrostatic steering, leveraging charge-based interactions to guide the correct assembly of heavy chains are based on the presence of oppositely charged amino acids. This approach can enhance antibody solubility, stability, or pharmacokinetics (41). Eight Fc variants for isoelectric point (pI) engineering, such as 14-G1v72 (negative) and 14-G1v73 (positive) in the CH3 domains of bsAb heavy chains. In our dataset, seven entries incorporate these variants, including *elranatamab, latikafusp, navicixizumab, petosemtamab*, *tepoditamab*, *zenocutuzumab,* and *besufetamig*.

Additionally, the controlled Fab-arm exchange (cFAE) method allows precise pairing of antibody arms by leveraging complementary interactions between the heavy chains, ensuring accurate bispecific antibody assembly (42). This strategy introduces a single mutation in the CH3 domain of each heavy chain: 18-G1v82-1 (CH3 L85.1) in one chain and 18-G1v82-2 (CH3 R88) in the other. Within IMGT/mAb-DB, this strategy has been identified in six entries.

Strategies to achieve correct light-heavy (H-L) pairing in bsAbs are less common than those targeting H-H heterodimerization, as engineering the heterodimeric CL/CH1 interface is more complex (43). Precise H-L pairing is critical since each heavy chain must correctly pair with its corresponding light chain to maintain antigen-binding specificity and functionality. The most widely used method, CrossMab® Technology (44), relies on domain swapping in the Fab region of one antibody arm. However, as this approach does not alter the AA sequence, it is excluded from our sequence-based analysis. Here, we focus on methods introducing complementary mutations in the constant regions of the light and heavy chains. These modifications promote selective interactions and ensure accurate H-L pairing.

We identified 21 Fc variants designed to optimize H-L pairing, present in 21 mAbs and one fusion protein (Table 1). The most frequently observed variants are 17-G1v57 (CH1 E26, E119) in the heavy chain and 17-KCv57 (C-KAPPA R12, K13) in the light chain (45). These modifications introduce complementary electrostatic and steric changes, creating a unique interaction surface that enhances selective H-L pairing.

**Table 1.**
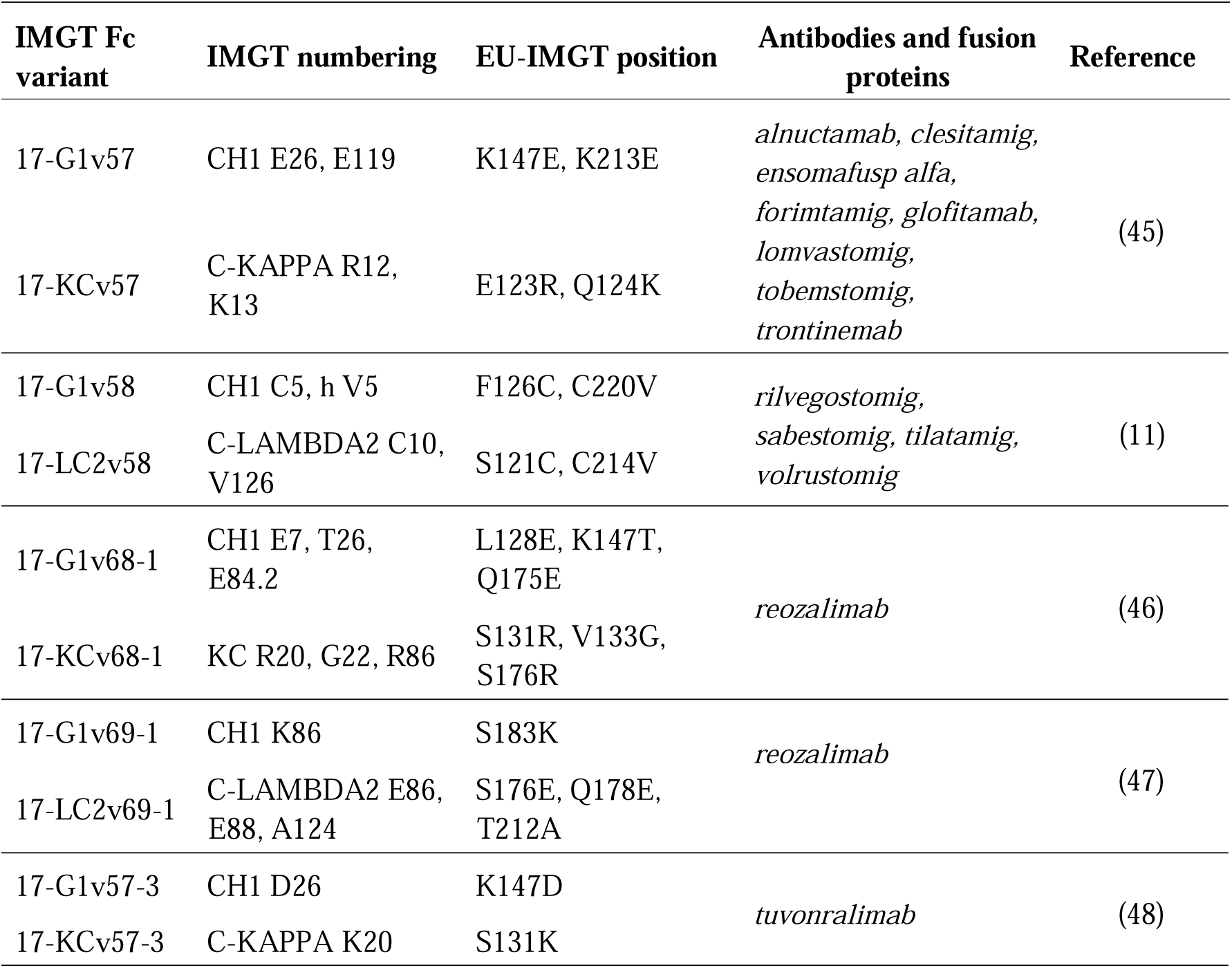
IMGT-described Fc variants involved in H-L chain heteropairing. For each variant, the IMGT numbering and EU-IMGT positions are provided, along with the antibodies and fusion proteins incorporating these modifications and relevant literature references.

*Reozalimab* (mAbID <u>1281</u>, LY3434172) presents an interesting design in bispecific antibody development. This antibody incorporates dual light chains, one kappa and one lambda, to ensure accurate H-L assembly. Specific mutations further enhance both H-L and H-H pairing. For kappa light chain-CH1 pairing, the 17-KCv68-1 in the C-KAPPA and 17-G1v68-1 in the CH1 are employed. Similarly, the lambda light chain-CH1 pairing utilizes the 17-LC2v69-1 variant in the C-LAMBDA2, paired with 17-G1v69-1 in CH1 partner (46). These modifications ensure optimal H-L pairing in *reozalimab*. For H-H pairing, *reozalimab* incorporates 14-G1v68 (CH3 V6, L22, L79) in one heavy chain and 14-G1v69 (CH3 V6, Y7, A85.1, V8) in the other. Additionally, to minimize its effector functions, *reozalimab* also includes 6-G1v14-67 (CH2 A1.3, A1.2, S27), which reduces ADCC and CDC activities (46,49). This antibody exemplifies the integration of multiple Fc engineering strategies, optimizing both structural and functional properties for therapeutic use.

In contrast, the 17-G1v57-3 and 17-KCv57-3 variants are not employed for bispecific formation but facilitate the production of antibodies mixtures within a single expression system, ensuring correct light chains pairing for *tuvonralimab* and *iparomlimab* (mAbID <u>1203</u>). In addition, *tuvonralimab* presents the 18-G1v82-7 variant in the CH3 domain (R84.2, E88) to control of H-H heteropairing, enhancing the fidelity of antibody assembly in the mixture (46).

Fc variants have also significantly advanced site-specific drug conjugation in ADCs. Most ADCs present in IMGT/mAb-DB lack engineered conjugation sites, but 32 ADCs carry specific Fc variants that introduce cysteine or other chemical handles at defined positions within the Fc region, enabling precise attachment of cytotoxic payloads while preserving antibody structure and function (50). Examples include 16-G1v27, 16-G1v44, and 16-G1v55, which introduce site-specific cysteines to improve ADC stability and homogeneity. The 16-G1v56 variant incorporates the non-natural amino acid *para*-azidomethyl-L-phenylalanine (pAMF), providing a precise site for covalent conjugation (51). These modifications support controlled drug-to-antibody ratio (DAR), enhancing the potency and safety of targeted therapies.

### ‘Physicochemical properties’ category

The ‘Physicochemical properties’ category focuses on Fc engineering strategies aimed at improving antibody stability, solubility, and manufacturability by modulating specific physicochemical characteristics (52). Key areas include enhancing protein and thermal stability, reducing aggregation, and optimizing the pI (53,54). Eighteen Fc variants were identified within this category, represented across 35 entries in IMGT/mAb-DB.

Protein stability is a fundamental determinant of therapeutic antibody’s ability to retain its functional conformation under stress conditions such as high temperatures, chemical exposure, or protease activity (52). Maintaining stability is therefore essential, as it directly affects both large-scale manufacturing and clinical performance. To address these challenges, Fc engineering strategies introduce targeted mutations within the Fc region that improve resistance to denaturation and unfolding. For example, the 10-G1v87 variant (CH2 V14, P90) confers thermal stability, whereas the 10-LC2v1 variant (C-LAMBDA2 G45, K82) enhances chain stability.

Aggregation is closely linked to protein stability and represents a critical challenge in antibody development, as it can compromise therapeutic efficacy, trigger immune-related side effects, and reduce bioavailability (55). Among IgG subclasses, IgG2 and IgG4 are more susceptible to aggregation than IgG1, particularly under conditions of low pH or heat-induced stress (56). To counteract this, Fc engineering strategies target specific AA within the Fc region to reduce aggregation propensity. Notable examples include the 6,10-G2v7 variant for IgG2 (57) and the 10-G4v8-1 variant for IgG4 (58), both of which mitigate acid-induced aggregation. These modifications help preserve antibodies in their monomeric, functional form during manufacturing, storage, and administration.

Another essential physicochemical parameter is the pI, which influences both solubility and stability of therapeutic antibodies. Antibodies with a more basic pI often demonstrate increased tissue uptake and faster clearance rates (59). Two Fc variants, 10-G1v88 (CH2 R94, CH3 R1.2) and 10-G1v88-1 (CH2 R94, CH3 K92), were identified as increasing antibody pI to enhance immune complexes. Beyond solubility and pharmacokinetics, pI engineering can also facilitate Fc heterodimer purification, as exemplified by the complementary variants 18-G1v82-5 (CH3 D24, S26) and 18-G1v82-6 (CH3 Q13, K20) (60).

Protein A is another important consideration in antibody engineering, as it influences purification efficiency and the avoidance of undesirable interactions during manufacturing. Fc engineered mAbs are often designed to abrogate protein A binding by introducing mutations that decrease affinity. For example, the variants 10-G4v8 (CH3 R115, F116, P125) and 10-G4v83 (CH3 R115, F116) specifically abrogate Protein A interaction, thereby facilitating controlled purification strategies.

Finally, Fc variants have been developed to eliminate immunogenic epitopes that may arise following directed mutagenesis. A notable case involves point mutation at the CH3 N114 (N434), introduced to prolong antibody half-life by enhancing FcRn affinity, this mutation was found to induce significant binding to rheumatoid factor (RF) (61), an autoantibody targeting the IgG Fc region (62). To reduce this, the 10-G1v89 variant (CH3 R118, E120) was introduced to disrupt the RF-binding site, maintaining its high affinity to FcRn (63). This strategy has been successfully applied in *crovalimab* (mAbID <u>783</u>), an anti-C5 mAb approved for the treatment of paroxysmal nocturnal hemoglobinuria.

Another immunogenicity-related innovation concerns aglycosylated IgG2 antibodies. AA changes in the conserved glycosylation site at CH2 84.4, by substituting asparagine with glutamine (N297Q) abolishes ADCC and CDC activities (8-G2v36 variant) but unexpectedly generates a novel T-cell epitope, thereby increasing immunogenicity. To overcome this, the 8,10-G2v36-104 variant introduces an additional substitution, a phenylalanine at position CH2 84.3 (F296A), which eliminates the newly formed T-cell epitope and reduces immunogenicity potential. This combined engineering strategy has been incorporated into *abituzumab* (mAbID <u>489</u>), an IgG2-IgG1 hybrid monoclonal antibody, thereby improving its immunological profile and enhancing therapeutic performance in oncology applications.

### ‘Hybrid’ category

Therapeutic antibodies are predominantly derived from the IgG1, IgG2, and IgG4 isotypes, each selected for their distinct structural and functional properties. Although all IgG subclasses share a long serum half-life, their immune effector functions differ: IgG1 is highly effective in complement activation and ADCC, IgG2 exhibits limited complement activity, and IgG4 generally lacks effector functions. In therapeutic contexts such as autoimmune diseases, minimizing effector function is crucial to reduce off-target effects and toxicity. A well-known example is *eculizumab* (mAbID <u>37</u>), which has been engineered to abrogate Fc-mediated activity. The ‘Hybrid’ category addresses these requirements that combine the advantageous features of different IgG subclasses, tailored to specific therapeutic needs.

Our dataset includes four hybrid combinations: IgG1-IgG2, IgG2-IgG1, IgG2-IgG4, and IgG4-IgG1. Among these, IgG1-IgG2 hybrid antibodies include *tafasitamab* (mAbID <u>522</u>) and *talacotuzumab* (mAbID <u>754</u>). These antibodies exhibit a unique structural configuration in which the CH1 domain, hinge region, and the N-terminus of the CH2 domain (lower hinge) are derived from the IGHG1*01 gene, while the CH2 and CH3 domains originate from the IGHG2*01gene, resulting in the Fc variant 19-G1G2v1. This hybrid structure enhances effector functions by combining the strong FcγR binding ability of IgG1 with the structural stability of IgG2. In addition, these antibodies incorporate the 2-G2v7-1 variant, which includes specific AA changes in the CH2 domain (D3, E117) to enhance ADCC. A further substitution, A110 (G327A), replaces an IgG2 residue with the corresponding IgG1 residue at the divergence point between the two isotypes (64,65).

In contrast, *abituzumab* primarily utilizes an IgG2 backbone but integrates the hinge region from the IGHG1*01 gene (19-G2G1v2 variant), conferring increased structural flexibility and stability (66). Similarly, *eculizumab* is an IgG2-IgG4 hybrid antibody (19-G2G4v1 variant) that combines the naturally low FcγR binding affinity of IgG2 with the absence of complement activation characteristic of IgG4. This dual attenuation of ADCC and CDC activities significantly reduces the risk of Fc-mediated adverse effects (67).

## Discussion

Several reviews have underscored the advances in Fc variant engineering (10,68), and initial analyses of engineered variants (21–23) have further highlighted the expanding repertoire of strategies available to tailor mAbs for specific therapeutic applications. Various databases, such as Thera-SAbDab (69) and Inxight Drugs (70), provide freely accessible data. In contrast, IMGT distinguishes itself by offering curated and standardized terminology based on IMGT-ONTOLOGY rules (17). IMGT/2Dstructure-DB integrates INN/WHO antibody-related sequences with detailed structural and gene-based annotation for describing immune receptors.

In this study, we demonstrate the utility of the IMGT/FcVariantsExplorer tool, which enables systematic analysis of Fc variants in therapeutic antibodies. By leveraging integrated data from IMGT/2Dstructure-DB, 1,310 monoclonal antibodies and fusion protein sequences were analysed, allowing precise identification and annotation of engineered variants. These variants were classified according to the standardized IMGT nomenclature for Fc-engineered variants (11,71).

Through our in-depth analysis of Fc-engineered antibodies, we identified 655 entries featuring modifications in the Fc region. These were classified into five functional categories: ‘Effector’, ‘Half-life’, ‘Physicochemical properties’, ‘Structure’, and ‘Hybrid’, which represent the broad range of engineering strategies employed to optimize mAb performance. In total, 251 unique Fc variants were annotated using the IMGT nomenclature for Fc variants. A comprehensive reference table is available on the IMGT website (https://www.imgt.org/IMGTbiotechnology/IGHG_variant/Tableau1.html). Interestingly, 44 entries in the dataset carry AA changes with unclear effects due to insufficient evidence in the literature to confirm their biological functions (Supplementary Table 1).

Fc engineering to modulate effector functions remains a central strategy in antibody design. Among the 336 mAbs with engineered Fc regions targeting effector functions, the vast majority (92%) were silenced to reduce immune activation, prioritizing safety by limiting effector responses, which is particularly relevant for autoimmune and inflammatory indications. Variants such as 6-G1v14 (LALA), which reduce ADCC and CDC through weakened FcγR and C1q binding (72), dominate this category. Nevertheless, the dataset also includes variants engineered to enhance effector functions, such as 2-G2v7-1 (73) and 2-G1v9-1 (74), which are of particular interest for oncology applications owing to their ability to improve ADCC. The 2-G1v9-1 variant incorporates a series of AA changes in the CH2 domain (L1.2>V, F7>L, R83>P, Y85.2>L) and one in the CH3 domain (P83>L), enhancing affinity for both allelic variants of FcγRIIIa and promoting a potent ADCC response against tumor cells, thereby contributing to strong anti-tumor activity (75).

Efforts to engineer antibody half-life have focused on modulating FcRn binding. Variants such as 9-G1v21 (CH2 Y15.1, T16, E18) can significantly extend mAb serum persistence by optimizing binding at acidic pH and release at neutral pH (76). Conversely, several antibodies in the dataset were deliberately modified to reduce FcRn binding, thereby shortening half-life as a therapeutic advantage in conditions where prolonged antibody persistence in the circulation could exacerbate disease, such as autoimmunity (77). However, the 9-G1v21 variant has been associated with reduced structural stability and an increased tendency for aggregation due to heightened local flexibility caused by AA changes in the CH2 domain (78). To overcome these limitations, the 6,9-G1v53 variant (CH2 L1.3>F, L1.2>Q, K105>Q) was developed, combining improved physiochemical properties while simultaneously reducing ADCC and CDC activities (79).

Structural stabilization also emerged as a major theme, underscoring the critical role of Fc modifications in improving antibody stability, particularly for IgG4 mAbs. Variants such as 12-G4v5, located in the hinge region, and 12-G4v6, in the CH3 domain, prevent Fab-arm exchange and preserve antibody integrity (37). These findings highlight the importance of Fc engineering in maintaining mAb structural stability. Additional approaches, including the “knob-into-hole” strategy and electrostatic steering, further enable heterodimerization for bispecific formats.

The cFAE method facilitates precise bispecific antibody heterodimerization by introducing single mutations in the CH3 domains (18-G1v82-1 and 18-G1v82-2 variants). These variants ensure accurate assembly and stability of the bsAb, are locked into their final configuration upon re-oxidation. Importantly, the cFAE method preserves the natural IgG structure, maintains effector functions, and ensures structural stability without compromising antibody solubility (42). As a result, bsAbs produced by this approach exhibit excellent pharmacokinetics, reduced aggregation, and optimized therapeutic activity, making the method suitable for clinical applications.

Fc engineering strategies aimed at optimizing physicochemical properties have also significantly advanced therapeutic antibody design. By enhancing stability, solubility, and pI while minimizing aggregation and immunogenicity, these variants ensure that antibodies meet clinical standards for both efficacy and safety. Improvements in thermal and chemical stability, such as those conferred by the 10-G1v87 and 10-LC2v1 variants, help maintain structural integrity under stress. Anti-aggregation mutations, including 6,10-G2v7 and 10-G4v8-1, further prevent functional loss during production and storage (57,58). Moreover, our dataset highlights the increasing application of hybrid variants, which combine structural elements from different IgG isotypes to harness the functional benefits of each across a wide range of disease conditions.

A major advancement presented in this study is the integration of Fc variant annotations into IMGT/mAb-DB. This enhancement enables users to perform targeted searches by variant function, ‘Category’ or ‘Effect’, thereby facilitating investigations of structure-function relationships and supporting the rational design of mAbs for specific clinical goals.

In conclusion, this study provides a comprehensive analysis of Fc-engineered monoclonal antibodies, offering curated annotations and functional classifications that reflect current strategies in therapeutic design. Through the integration of variant data into IMGT/mAb-DB and the use of the IMGT/FcVariantsExplorer tool, researchers can explore Fc mutations with both scale and precision. Continued updates aligned with INN/WHO lists and regulatory data from FDA and EMA, together with standardized variant annotations according to the IMGT nomenclature, will further reinforce IMGT’s role as pivotal resource for antibody engineering and design.

## Methods

### IMGT/2Dstructure-DB dataset

The analysed dataset comprises AA sequences from 1,482 antibodies and fusion proteins, available in IMGT/2Dstructure-DB. These sequences are derived from the INN/WHO Proposed and Recommended Lists (13) and have been annotated according to the IMGT Scientific Chart rules (17,18). Annotations follows the standardized axioms of IDENTIFICATION (keywords), DESCRIPTION (labels in capital letters), CLASSIFICATION (gene and allele names) and NUMEROTATION (IMGT unique numbering and IMGT Collier de Perles for V domain (19) and for C domain (20)).

### IMGT® nomenclature of engineered Fc variants

The IMGT® nomenclature of engineered Fc variants characterizes Fc variants involved in antibody effector properties and formats (11,71). This nomenclature also applies to antibody chains other than the gamma isotype, such as light chains (kappa or lambda) and the joining chain, as well as to species other than *Homo sapiens* (e.g., *Mus musculus*, *Canis lupus familiaris*) (11,12). In this study, the IMGT® nomenclature of engineered variants has been extended to include a new category, ‘Hybrid’, which groups mAbs and fusion proteins containing constant domains from different isotypes. Thus, the IMGT® nomenclature now classifies 19 variant types grouped into five categories: ‘Effector’ (ADCC, ADCP and CDC), ‘Half-life’, ‘Physicochemical properties’, ‘Structure’, and ‘Hybrid’ (Table 2).

**Table 2.**
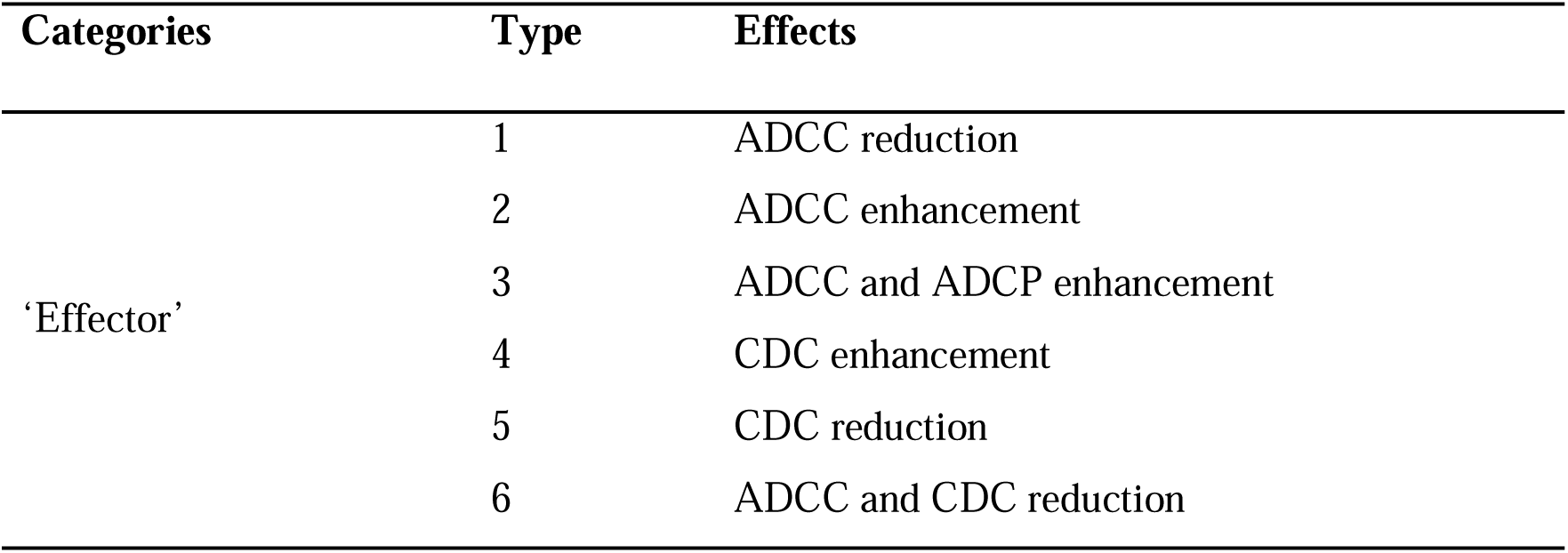
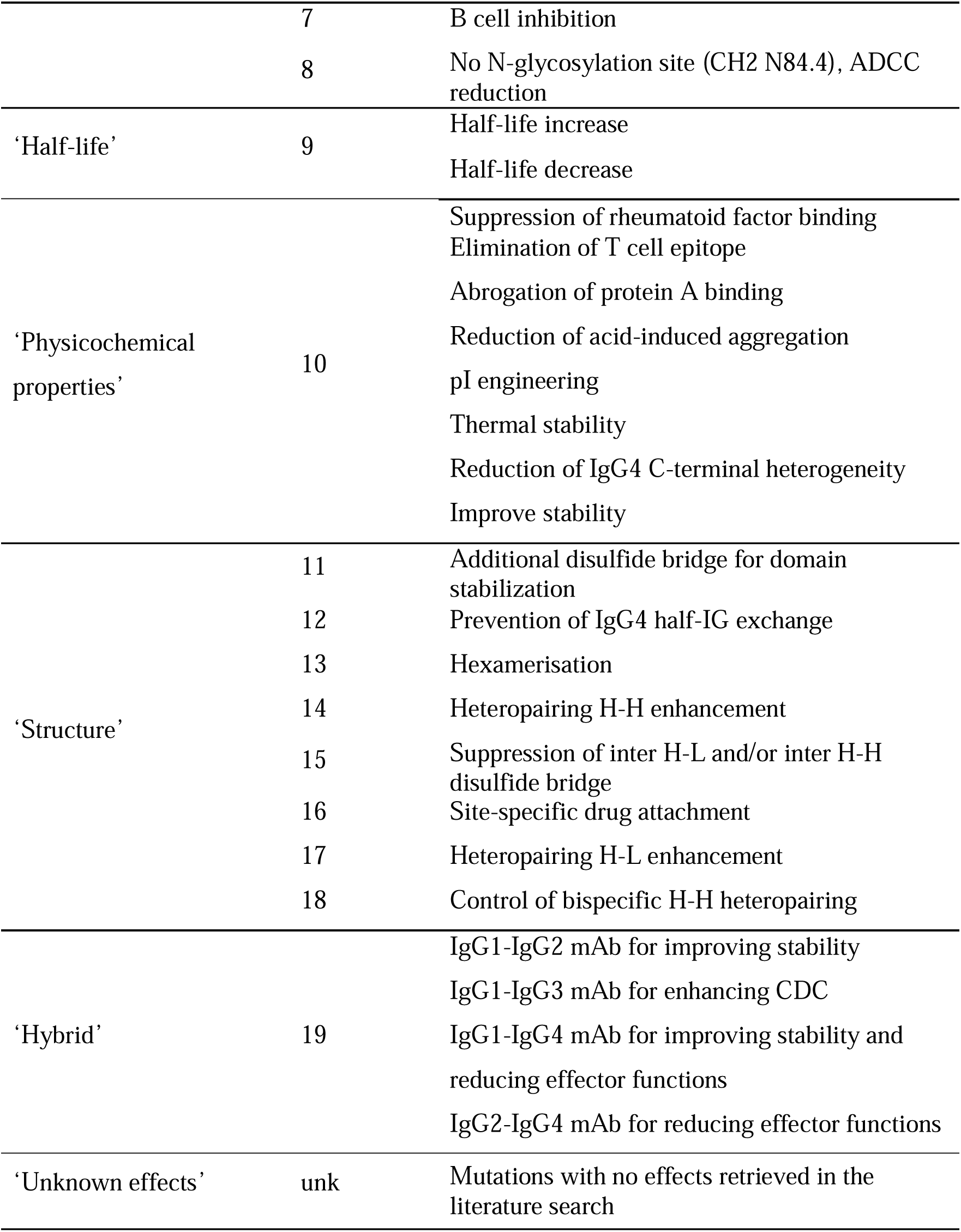
IMGT® nomenclature of engineered Fc variants classified into five categories, comprising 19 types based on their distinct biological effects (11).

An IMGT-engineered Fc variant name comprises the species (abbreviated), one or several numbers from 1 to 19 indicating the property and function type(s) (Table 1), followed by a dash, the gene name (abbreviated), the letter ‘v’, and a unique number (e.g., Homsap 1-G1v1) (11). The IMGT variant definition also specifies, per domain (e.g., CH2), the amino acid(s) of the variant using the one-letter abbreviation along with its position according to the IMGT unique numbering for C domains (19); for example: Homsap 1-G1v1 CH2 P1.4 (11). The Eu-IMGT positions (80) have been defined and formalized to enable comparisons with other studies.

### Data processing

To perform a comprehensive analysis of the dataset, we focused on the AA sequences from 1,216 antibodies chains and 94 fusion proteins chains that include at least one constant domain (CH1, HINGE-REGION, CH2, CH3 for heavy chains; C-KAPPA, C-LAMBDA for light chains). We developed the IMGT/FcVariantsExplorer tool, an automated tool designed to detect AA changes by comparing the sequences with the IMGT reference directory (14). The interface web of IMGT/FcVariantsExplorer allows the user to upload and submit AA sequences from IG or FPIA chains. Each domain sequence is aligned with the corresponding domain of the closest gene and allele, and the tool automatically records AA change data. In addition to substitutions, IMGT/FcVariantsExplorer identifies insertions and deletions by directly comparing aligned AA sequences, thereby accurately pinpointing these variations and their positions (Figure 4).

**Figure 4.**
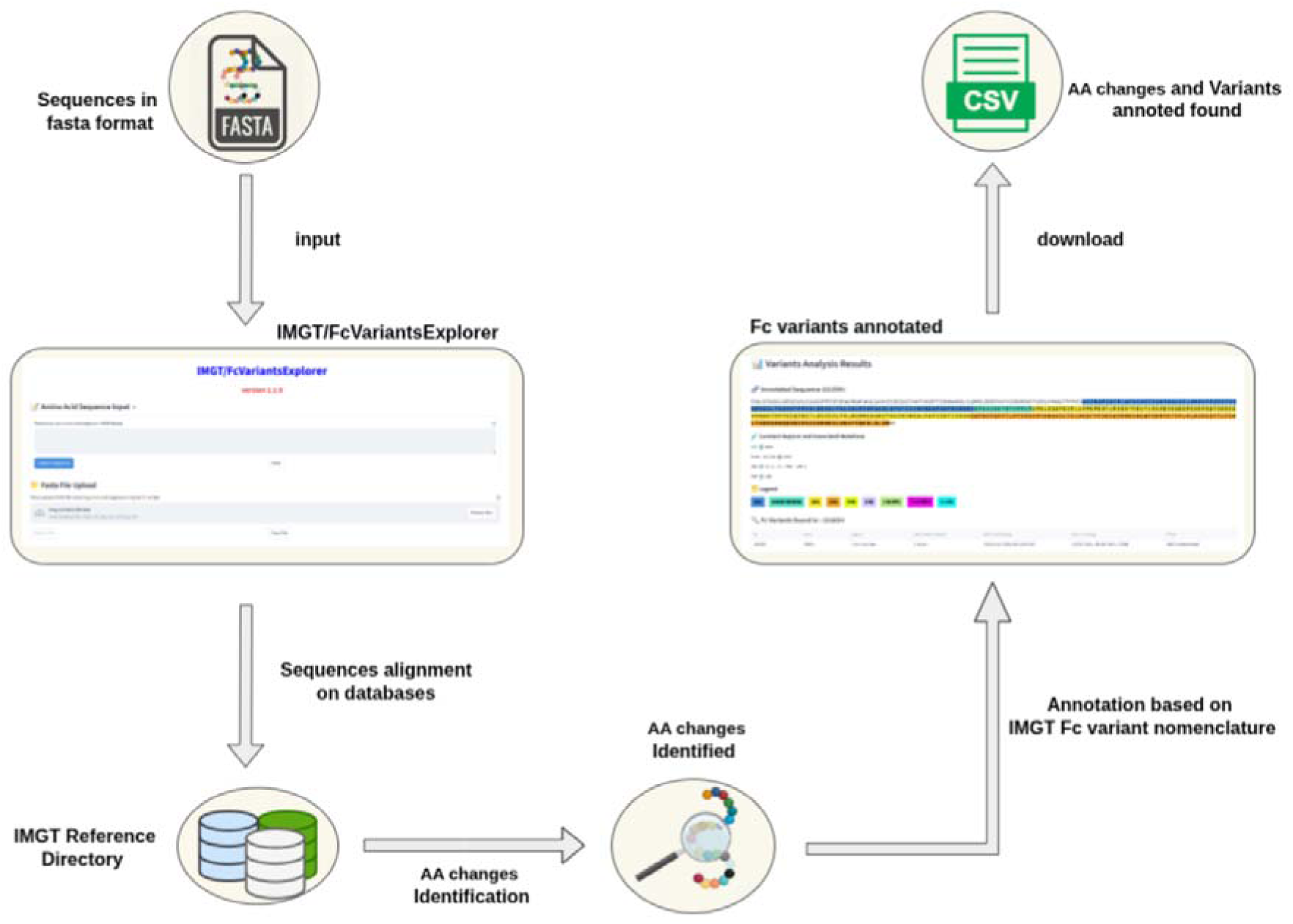
Flowchart illustrating the *IMGT/FcVariantsExplorer* workflow. The process begins with an input amino sequence in FASTA format. The tool aligns this sequence against the IMGT reference directory, after which AA changes are identified. These changes are then cross-referenced with Fc variants already annotated by IMGT to assign the corresponding Fc variant nomenclature. The analysis output is provided as CSV (Comma-Separated Value) file containing the detected AA changes and their annotated variants.

Once AA changes are annotated, the tool determines the engineered variant through a structured matching process. First, the detected AA changes and identified isotype are compared with the IMGT® nomenclature of engineered variants. If no exact match is found, a secondary matching process is triggered to explore potential combinations of AA changes present in the sequence. This approach enables a more comprehensive identification of possible variants.

### Literature mining

To designate nomenclature for new Fc variants, we developed an *in-house* text mining tool for literature searches. The main goal of this tool is to identify and classify the properties and functions associated with Fc region variation, organizing them according to variant categories and their respective effects. Amino acid changes detected by the IMGT/FcVariantsExplorer tool that do not correspond to any known Fc variants are further investigated using this automated approach to assess their potential biological functions.

Briefly, the tool employs a text-mining strategy to select, classify, and analyse a diverse range of textual sources, including PubMed, PubMed Central (PMC) Open Access full-text articles. Through this comprehensive search process, it extracts critical biological effects associated with Fc variants.

The text mining pipeline was implemented in Python and is based on a dictionary matching approach, enhanced with parallel multiprocessing to efficiently screen articles. The dictionary includes identified AA changes, enabling the tool to extract relevant information regarding their biological effects (Figure 5). The extracted data were subsequently curated manually to ensure accuracy and to interpret the results of the text mining process. This methodology enabled us to establish the biological effects of newly identified AA changes in antibody sequences, with each IMGT nomenclature for engineered variant supported by at least one literature reference.

**Figure 5.**
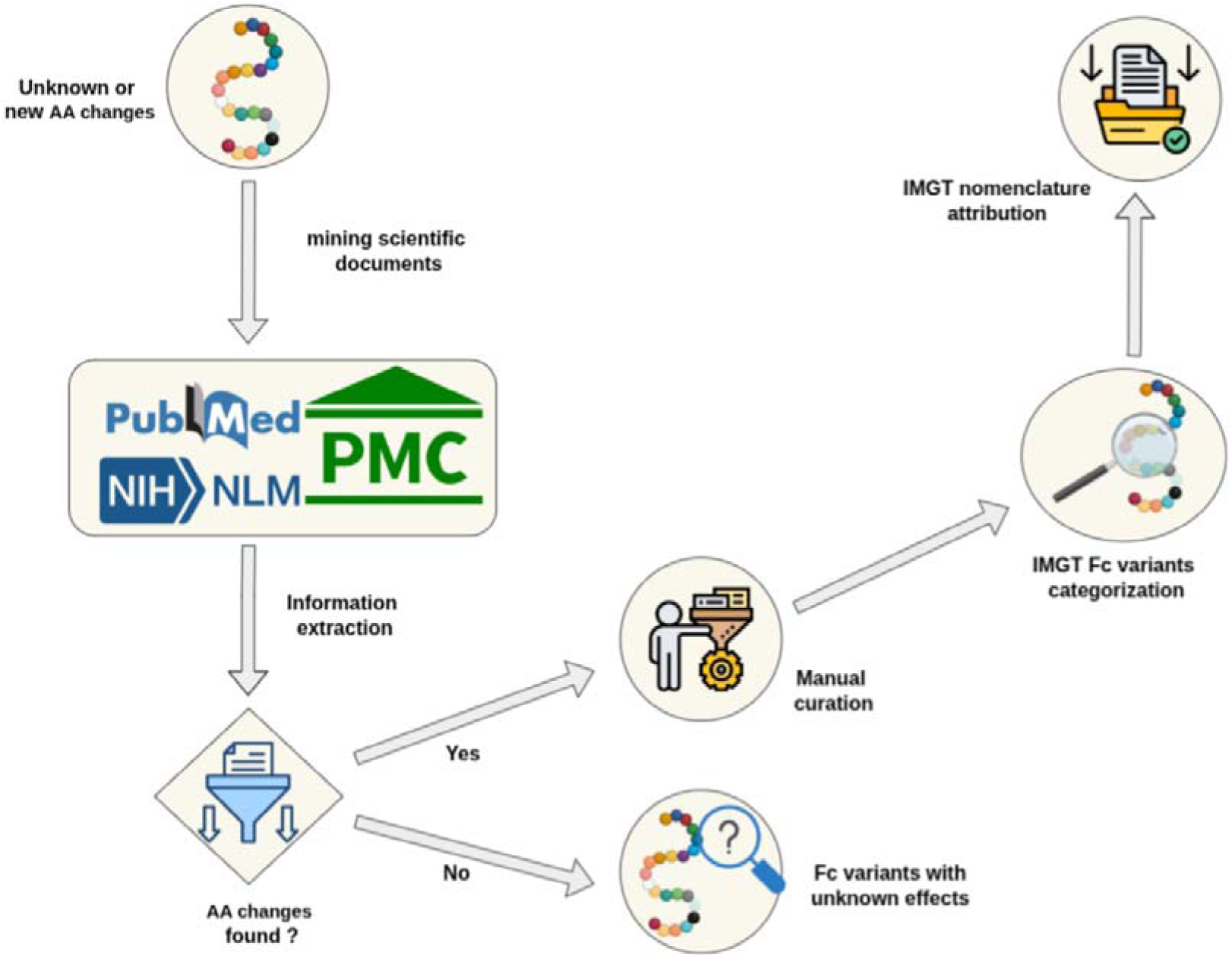
Amino acid changes in our dataset that do not correspond to Fc variants annotated by IMGT are subjected to a literature search using an in-house text mining and pattern-matching tool. This tool extracts and classifies the biological effects from indexed scientifical articles. The results are then manually curated to ensure accuracy and to assign new Fc variant nomenclature. Variants for which no biological effects are identified are labeled as ‘unknown effects’. To keep the database current, literature mining is performed monthly to capture newly published studies related to these Fc variants.

This integrative approach, combining automated literature mining with expert manual curation, ensures scalable and reliable annotation of Fc-engineered variants. By systematically linking each variant to published experimental data, we provide a robust and transparent framework for updating IMGT nomenclature. Ongoing developments at IMGT aim to enhance the IMGT/FcVariantsExplorer tool by integrating the literature mining pipeline into a new user-accessible version.

## Acknowledgements

We thank all members of the IMGT® team for their expertise and constant motivation. We are grateful to Gérard Lefranc for his contribution to the antibody engineered variants nomenclature. We acknowledge the support of Immun4Cure IHU “Institute for innovative immunotherapies in autoimmune diseases” (France 2030/ANR-23-IHUA-0009). IMGT® is member of the French Infrastructure Institut Français de Bioinformatique (IFB) as well as member of BioCampus, MAbImprove and IBiSA.

## Data availability statement

The datasets analyzed in the current study are available in IMGT/mAb-DB and in IMGT/2Dstructure-DB of IMGT® the International ImMunoGeneTics Information System®.

## Authors’ contributions

TM conceived, analyzed the data, and drafted the manuscript. CN, IM, and GS developed the tool. FB supervised the text-mining approach. VG discussed and drafted the manuscript. PD analyzed the data and developed the databases. MPL conceived the variant nomenclature, analyzed the data and the findings. SK conceived and supervised the findings and the write up of this work. All authors contributed to the article and approved the submitted version.

## Abbreviations

3D: three-dimensional
AA: amino acid
ADCC: antibody-dependent cellular cytotoxicity
ADCP: antibody-dependent cellular phagocytosis
CDC: complement-dependent cytotoxicity
CPCA: composite protein for clinical application
EMA: European Medicines Agency
Fc: Fragment crystallizable
FcRn: neonatal Fc receptor
FcγR: Fc gamma receptors
FDA: Food and Drug Administration
FPIA: fusion protein for immune application
INN: International Nonproprietary Name
mAb: monoclonal antibody
PMC: PubMed Central
RPI: related protein of the immune system
TR: T cell receptor
WHO: World Health Organization.

---

**Figure 6.** Extract from the IMGT/mAb-DB results page illustrating a query to retrieve mAbs or fusion proteins engineered to silence Fc effector properties. The search combines the filters ‘Effect: ADCC and CDC reduction’ and ‘Clinical domain: Immunology’, resulting in the identification of 49 molecules specifically designed to minimize Fc-mediated effector activity in the context of autoimmune diseases.

**Figure 7.** Distribution of Fc-engineered monoclonal antibodies across the five IMGT variant categories: ‘Effector’, ‘Half-life’, ‘Physicochemical properties’, ‘Structure’, and ‘Hybrid’. The counts represent the number of molecules carrying at least one Fc variant within each category. As individual antibodies may include multiple Fc variants, a single molecule can be represented in more than one category.

**Figure 8.** Frequency of the three most common ‘Effector’ Fc variants across IgG subclasses. IgG1 variants are represented in blue, IgG2 in yellow, and IgG4 in green.

**Figure 9.** Flowchart illustrating the *IMGT/FcVariantsExplorer* workflow. The process begins with an input amino sequence in FASTA format. The tool aligns this sequence against the IMGT reference directory, after which AA changes are identified. These changes are then cross-referenced with Fc variants already annotated by IMGT to assign the corresponding Fc variant nomenclature. The analysis output is provided as CSV (Comma-Separated Value) file containing the detected AA changes and their annotated variants.

**Figure 10.** Amino acid changes in our dataset that do not correspond to Fc variants annotated by IMGT are subjected to a literature search using an in-house text mining and pattern-matching tool. This tool extracts and classifies the biological effects from indexed scientifical articles. The results are then manually curated to ensure accuracy and to assign new Fc variant nomenclature. Variants for which no biological effects are identified are labeled as ‘unknown effects’. To keep the database current, literature mining is performed monthly to capture newly published studies related to these Fc variants.

