## Supplemental Table 1 for "Identification of Engineered IMGT Fc Variants in IMGT/mAb-DB Therapeutic Antibodies and Fusion proteins"

**Supplementary Table 1**. Fc variants with unknown effects in engineered monoclonal antibodies. Species, antibody, provisional variant nomenclature, and amino acid changes (IMGT and EU numbering) are shown. Probable functional effects are indicated when the mutations can be related to existing Fc variants.

| Specie | Molecule name | Allele | Variant nomenclature | IMGT numbering | EU numbering | Probably effects |
|---|---|---|---|---|---|---|
| *Homo sapiens* | anetumab, teclistamab, xentuzumab | IGLC2*01 | unk-1 | LC G43 | A150G | |
| | bavituximab | IGKC*01 | unk-2 | KC A1.3 D1.2 | T109A, V110D | |
| | canvircept | IGHG1*01 | unk-3 | CH3 T125 | P445T | |
| | cinpanemab | IGLC2*01 | unk-4 | LC S1.5 | G107S | |
| | cixutumumab, lodapolimab | IGLC2*01 | unk-5 | LC A124 | T212A | |
| | conbercept | IGHG1*01 | unk-6 | h L13, CH3 A80 | P228L, T393A | ADCC enhancement |
| | derlotuximab, frexalimab, galiximab | IGHG1*01 | unk-7 | CH1 A121 | V215A | ADCC and CDC reduction |
| | galiximab | IGLC2*01 | unk-8 | LC2 T3 | S114T | |
| | efaprinermin alfa | IGHG1*03 | unk-9 | CH3 K15, T16 | T359K, K360T | |
| | efdelikofusp alfa, efzilonkofusp alfa | IGHG4*01 | unk-10 | h ^1 ins^A | A215^216 | |
| | efineptakin alfa, efmedaglutide alfa | IGHD*01-IGHG4*01 | unk-11 | IGHD*01 (h-CH2)<br>IGHG4*01 (CH2-CH3) | | |
| | efdoralprin alfa | IGHG4*01 | unk-12 | CH2 Y15.1, CH3 L107 | M252Y, M428L | Half-Life increase |
| | efgivanermin alfa | IGHG1*03 | unk-13 | h L5 | C220L | Inter H-L disulfide bridge suppression |
| | ensituximab | IGHG1*01 | unk-14 | CH1 P85.3 | L179P | |
| | eptinezumab | IGHG1*03 | unk-15 | CH1 A119 | K213A | |
| | iratumumab | IGHG1*01 | unk-16 | CH3 delY116 | Y436del | Half-life increase<br>Protein A binding abrogation |
| | lesigercept | IGHD*01 | unk-17 | IgD h G43, S44 | K192G, E193S | |
| | lucatumumab | IGHG1*03 | unk-18 | CH1 A10 | S131A | |

| Specie | Molecule name | Allele | Variant nomenclature | IMGT numbering | EU numbering | Probably effects |
|---|---|---|---|---|---|---|
| | metelimumab | IGHG4*01 | unk-19 | CH2 P13 | D249P | |
| | metelimumab | IGKC*01 | unk-20 | KC delS114 | S202del | |
| | nemolizumab, satralizumab | IGHG2*01 | unk-21 | CH1 S10, K12; CH2 Q30 | C131S, R133K, H268Q | ADCC and CDC reduction |
| | nemolizumab | IGHG4*01 | unk-22 | CH3 M84 | V397M | |
| | necitumumab, ramucirumab | IGHG1*03 | unk-23 | CH1 L5 | F126L | |
| | nimotuzumab | IGKC*01 | unk-24 | KC E1.3 | T109E | |
| | olinvacimab, ulenistamab | IGKC*01 | unk-25 | KC S1.3 | T109S | |
| | ontuxizumab | IGHG1*01 | unk-26 | CH3 F85.3 | S403F | B cell inhibition |
| | opucolimab | IGLC3*03 | unk-27 | LC delG1.5, R119, L123, delS127 | G107Adel, K207R, P211L, S215del | |
| | ormutivimab | IGHG1*03 | unk-28 | CH1 L3 | S124L | |
| | otelixizumab | IGLC2*01 | unk-29 | LC2 R1.5 | G107R | |
| | prafnosbart | IGKC*01 | unk-30 | KC A1.3 | T109A | |
| | ramucirumab | IGKC*01 | unk-32 | KC G1.4 | R108G | |
| | satralizumab | IGHG4*03 | unk-33 | CH3 M84, A114 | V397M, N434A | Half-life increase |
| | satumomab | IGKC*01 | unk-34 | KC A1.3, D1.2, T3, delR123, delG124, delE125, delC126 | T109A, V110D, S114T, R211del, G212del, E213del, C214del | |
| | satumomab | IGHG4*01-IGHG2A*01 | unk-35 | IGHG4*01 (CH1) human IGHG2A*01 (CH2-CH3) mouse | | |
| | silevimig | IGHG1*03 | unk-36 | CH3 G84.4, D85.4 | D401G, G402D | |
| | telisotuzumab | IGHG1*03 | unk-37 | h C7, delT8, delT10 | K222C, T223del, T225del | |
| | tividenofusp alfa | IGHG1*01 | unk-38 | CH3 Y44, delQ45.1, T45.2, W45.4, 45.7ins^A, T92, E94, E95, F100 | N384Y, Q386del, P387T, N389W, A389^390, D413T, S415E, R416E, N421F | |
| | urabrelimab | IGKC*01 | unk-39 | KC delR1.4 | R108del | |

| Specie | Molecule name | Allele | Variant nomenclature | IMGT numbering | EU numbering | Probably effects |
|---|---|---|---|---|---|---|
| | ustekinumab | IGHG1*01 | unk-40 | CH1 S1.4 | A118S | |
| | visilizumab | IGHG2*01 | unk-41 | CHS S1 | G446S | |
| *Felis catus* | dovanvetmab | IGHG1*01 | Felcat unk-1 | CH2 A1.3, A1.2, A1 | M234A, L235A, G237A | ADCC and CDC reduction |
| | dovanvetmab | IGKC*01 | Felcat unk-2 | KC Q122, delQ127, delR128, delE129 | N210Q, Q215del, R216del, E217del | |
| | relfovetmab | IGKC*01 | Felcat unk-3 | KC Q122 | N210Q | |
| *Canis lupus familiares* | blontuvetmab | IGHG2*02 | Canlupfam unk-1 | CH1 Q14 | T135Q | |
| | gilvetmab | IGKC*01 | Canlupfam unk-2 | KC ^S84.3 | S167^168 | |
| | riltovetbart | IGHG2*02 | Canlupfam unk-3 | CH3 H114 | N434H | |
| *Mus musculus* | apamistamab | IGHG1*02 | Musmus unk-2 | CH1 E100; CH2 Q81, delS85, F84.3; CH3 D27 | Q196E, K290Q, I296F, S302del, N371D | |
| | lemalesomab, racotumomab | IGHG1*01 | Musmus unk-3 | CH1 Q84.2 | E175Q | |
| | lilotomab | IGHG1*01 | Musmus unk-4 | CH1 Q84.2, T95, W96; CH3 D84.2, D84.4 | E175Q, P192T, R193W, N399D, N401D | |
| | miromavimab, omburtamab | IGHG1*02 | Musmus unk-5 | CH1 E100; CH2 Q81, F84.3; CH3 D27 | Q196E, K290Q, I296F, N371D | |
| | muromonab-CD3 | IGHG2A*01 | Musmus unk-6 | CH1 G13 | D134G | |
| | muromonab-CD3 | IGKC*01 | Musmus unk-7 | C-KAPPA T1.1 | A111T | |
| | nacolomab | IGHG1*02 | Musmus unk-7 | CH1 E100 | Q196E | |
